# Pesticide-induced Alterations to Phytoplankton Abundance and Community Structure Alter Ecosystem Respiration: Implications for the Carbon Cycle?

**DOI:** 10.1101/2020.10.15.341065

**Authors:** Samantha L. Rumschlag, Dale A. Casamatta, Michael B. Mahon, Jason T. Hoverman, Thomas R. Raffel, Hunter J. Carrick, Peter J. Hudson, Jason R. Rohr

**Affiliations:** Department of Biological Sciences, Eck Institute for Global Health, and Environmental Change Initiative, University of Notre Dame, Notre Dame, IN, USA; Department of Biology, University of North Florida, Jacksonville, FL, USA; Department of Forestry and Natural Resources, Purdue University, West Lafayette, IN, USA; Department of Biological Sciences, Oakland University, Rochester, MI, USA; Department of Biology and Institute for Great Lakes Research, Central Michigan University, Mount Pleasant, MI, USA; Department of Biology Pennsylvania State University, State College, PA, USA

**Keywords:** aquatic ecology, pesticides, synthetic chemicals, phytoplankton, zooplankton, carbon cycle, ecosystem respiration, disturbance

## Abstract

Current predictions of the effects of synthetic chemicals on freshwater ecosystems are hampered by the sheer number of chemical contaminants entering aquatic systems, the diversity of organisms inhabiting these systems, and uncertainties about how contaminants alter ecosystem metabolism. We conducted a mesocosm experiment that elucidated the responses of ponds composed of phytoplankton and zooplankton to standardized concentrations of 12 pesticides, nested within four pesticide classes and two pesticide types. We show that the effects of the pesticides on algae were consistent within herbicides and insecticides and responses of over 70 phytoplankton species and genera were consistent within broad taxonomic groups. Insecticides generated top-down effects on phytoplankton community composition and abundance, which were associated with persistent increases in ecosystem respiration. Herbicides reduced phytoplankton abundance, which was associated with decreases in primary productivity and ecosystem respiration. These results suggest that widespread pesticide use could have underexplored implications for the global carbon cycle. While these effects on ecosystem respiration were mediated through complex effects on communities, taxonomic groups of organisms responded similarly to pesticide types, suggesting opportunities to simplify ecological risk assessment.

## Introduction

The importance of aquatic systems to life on earth cannot be overstated. Humanity relies on freshwater as a precious natural resource for drinking, food production, and carbon capture (Meybeck 2003, Vörösmarty et al. 2005, World Water Assessment Programme 2009). Freshwater systems are hotspots for biodiversity (Dudgeon *et al.* 2006; Balian *et al.* 2008), and they play a crucial role in the capture, storage, and release of atmospheric carbon (Holgerson & Raymond 2016). Yet, the benefits of economic productivity, spurred by access to water, have been accompanied by impairments that threaten aquatic systems (Vörösmarty *et al.* 2010). In particular, the increasing rate of synthetic chemical pollution, which outpaces all other global change drivers, presents a major threat to freshwater systems globally (Bernhardt *et al.* 2017). In the United States alone, more than 500 million pounds of pesticide active ingredients are applied every year (Atwood & Paisley-Jones 2017). These applications have led to well-documented and widespread contamination of freshwater (Dudgeon *et al.* 2006; Gilliom & Hamilton 2006; Stone *et al.* 2014) and are implicated as a contributing factor to biodiversity declines in these systems (Malaj *et al.* 2014; Stehle & Schulz 2015).

Despite the magnitude of the threat that pesticide pollution imposes on freshwater systems, it is somewhat surprising that ecologists and their funding agencies have largely ignored the intricacies that these disturbances can yield on communities and ecosystems (Bernhardt *et al.* 2017; Burton *et al.* 2017). For instance, while pesticides have the potential to alter carbon cycling in freshwater systems via changes in the abundance or composition of community members, few studies have evaluated this possibility (but see McMahon *et al.* 2012; Halstead *et al.* 2014; Rumschlag *et al.* 2020). Classic toxicological research has employed lab-based exposure experiments, documenting that scores of pesticides harm model organisms. While these tests have been crucial in uncovering mechanisms of toxicity and are the basis of chemical regulation around the world, they often fail to predict the complex suite of effects that occur when pesticides enter freshwater systems (Gessner & Tlili 2016; Rohr *et al.* 2016; Bernhardt *et al.* 2017). For instance, traditional toxicological tests that focus on single pesticides and single organisms under controlled conditions ignore indirect effects of pesticides mediated through species interactions and food web structures (Kidd *et al.* 2014; Rumschlag *et al.* 2019). Instead, if the focus is on assessing the safety of pesticides on ecological systems, more realistic experimental conditions, like those of replicated field-based mesocosm experiments, should be incorporated into a framework of risk assessment. Mesocosms mimic well the complexity of multitrophic communities, so that both direct and indirect effects of pesticides can be evaluated (Rohr *et al.* 2006; Clements & Rohr 2009). In addition, testing that incorporates naturally complex communities allows for the evaluation of whether pesticide exposure can directly or indirectly alter ecosystem functions, including ecosystem metabolism, which is not possible with the traditional single species tests (Bernhardt *et al.* 2010; Halstead *et al.* 2014; Gessner & Tlili 2016).

In the U.S. and Europe, tens of thousands of synthetic chemicals are registered for use, and in the U.S. alone more than 350 pesticides are applied annually in agriculture (Baker & Stone 2015). Further, freshwater systems are home to about 125,000 described species, even though they only occupy 0.8% of earth’s surface (Dudgeon et al. 2006, Balian et al. 2008). Predicting the cumulative effects of pesticides on freshwater systems is enormously challenging because of the diverse array of pesticides to which ecosystems are exposed, combined with the diversity of organisms that exist in freshwater systems. Predicting these effects could be simplified if the effects of pesticides are similar within pesticide types (e.g. insecticides and herbicides that are designed to target insect and plant pests, respectively) or pesticide classes (i.e. chemical classes of pesticides that share similar chemical structures and molecular targets within a pest). Further, predicting the effects of pesticides could be simplified if organisms that are taxonomically related or share similar functional roles within an ecosystem have similarities in their responses to pesticides, a trend that has been shown in previous toxicological research (Ippolito *et al.* 2012; Hua & Relyea 2014). For example, our previous research has shown consistency in the effects of pesticides by class and type on parasite transmission, ecosystem functions, and macroinvertebrate and amphibian communities (Rumschlag *et al.* 2019, 2020). In addition, organisms that share functional roles in a community have been shown to respond similarly to types and classes of pesticides (Rumschlag *et al.* 2020). But to date, no study has attempted of evaluate the consistency of responses of phytoplankton community members to pesticide classes and types.

In the current study, we conducted an outdoor, replicated mesocosm experiment focused on exploring the effects of a diverse array of pesticides on phytoplankton and zooplankton communities and their contributions to ecosystem respiration. Phytoplankton and zooplankton were chosen as focal communities because: both groups have relatively short generation times (e.g. upwards of a day for phytoplankton (Laws 2013) and weeks for zooplankton (Kalff 2002)) allowing them to establish population dynamics within the duration of this experiment, both groups are taxonomically diverse allowing for the assembly of diverse communities in this experiment, and finally both groups are important contributors to ecosystem metabolism (Kalff 2002). Algae and macrophytes convert sunlight into biomass via photosynthesis, and zooplankton are the primary consumers of algae. Our objectives were to: 1) determine the consistency in the effects of pesticides by type, class, and individual pesticide on freshwater phytoplankton communities, 2) gauge the consistency of responses of phytoplankton community members to pesticides within five broad taxonomic groups (green algae, diatoms, cryptophytes, cyanobacteria, and euglenoids), 3) evaluate how herbicides and insecticides alter phytoplankton abundance and community composition via direct toxicity and indirect, top-down effects mediated by the zooplankton community, and 4) examine how respiration of aquatic systems is altered by pesticide-induced perturbations to aquatic communities over time.

We proposed four hypotheses. First, we hypothesized that the effects of pesticides would be consistent within pesticide types and classes; pesticides with similar taxonomic targets or similar chemical structures would have similar effects on phytoplankton communities. This hypothesis is motivated by the nestedness of the biologic activity of pesticides by class and by type. Pesticides of similar types (e.g. insecticides, herbicides) have similar targets in the environment (e.g. insects and plants, respectively), so pesticides within types likely have similar effects on non-target taxa (e.g. zooplankton and algae, respectively). Classes of pesticides share modes of action, meaning that they target the same biochemical and molecular pathways (e.g. triazine herbicides bind to the QB protein in the photosystem II reaction center blocking photosynthesis), which would drive similarity of observed effects within classes on phytoplankton communities. Second, we hypothesized that the responses of phytoplankton to pesticide exposures would be similar within broad taxonomic groups; a trend shown in other studies (Peterson *et al.* 1994; Ippolito *et al.* 2012; Hua & Relyea 2014). Third, we hypothesized that herbicides would cause direct reductions in abundance of phytoplankton across broad taxonomic groups, while insecticides would increase phytoplankton abundance and alter community composition through top-down effects on the zooplankton community. And finally, we hypothesized that pesticides would induce changes in ecosystem respiration; the duration of these changes would be explained by the environmental persistence of the pesticides and the generation time of the organism to which the pesticide is directly toxic. For instance, we predicted that respiration in communities exposed to herbicides might recover quickly because photosynthetic phytoplankton have short generation times and thus might rebound rapidly from the direct toxicity of herbicides. In contrast, respiration in communities exposed to insecticides might recover more slowly because direct toxicity occurs to longer-lived zooplankton that might have top-down effects on algae. In addition, we predicted that pesticides that persist for short durations of time in the environment would cause short-term disruptions to respiration because as the pesticide degrades, the community would recover more quickly from the initial perturbation relative to more persistent pesticides.

## Methods

### Aquatic Communities and Experimental Design

A randomized-block experiment was performed at the Russell E. Larsen Agricultural Research Center (Pennsylvania Furnace, PA, USA) using replicated mesocosm ponds. Mesocosms were 1,100 L cattle tanks covered with 60% shade cloth lids. The spatial block was distance from a tree line in the mesocosm field. We filled mesocosms with 800 L of water and 300 g mixed hardwood leaves. Mesocosms were inoculated with zooplankton, phytoplankton, and periphyton that were homogenized from four local ponds. Three weeks later, after these additions, pesticides were applied. To mimic the complex food web structure of natural ponds, we also added two snail, three larval anuran, one larval dragonfly, one water bug, one water beetle, one larval salamander, and one backswimmer species to each mesocosm on the day of pesticide applications, just prior to application. More specifically, each mesocosm received 11 *Helisoma (Planorbella) trivolvis*, 10 *Physa gyrina*, 20 *Hyla versicolor*, 20 *Lithobates palustris*, 20 *Lithobates clamitans*, 2 *Anax junius*, 2 *Belostoma flumineum*, 5 *Hydrochara* sp., 3 *Ambystoma maculatum*, and 6 *Nototeca undulata.* Responses of these community members to the established treatments are explored in Rumschlag *et al.* 2020 and are not a focus of the current study.

We randomly assigned 14 treatments (12 pesticides, 2 controls) with four replicate mesocosms of each treatment, which resulted in 56 total mesocosms (Fig. 1A). The 12 pesticide treatments were nested; treatments included two pesticide types (insecticide, herbicide), two classes within each pesticide type (organophosphate insecticide, carbamate insecticide, chloroacetanilide herbicide, triazine herbicide), and three different pesticides in each of four classes (Fig. 1A). Samples from two mesocosms were not processed for phytoplankton identification because of an error, so all analyses contain only 54 total mesocosms with three replicates for acetochlor and simazine treatments. At the start of the experiment, we applied a single dose of technical grade pesticides at environmentally relevant concentrations to mimic runoff of pesticides into freshwater systems following rainfall. To calculate environmentally relevant concentrations, we used U.S. Environmental Protection Agency’s GENEEC v2 software to generate estimated environmental concentrations of pesticides. We acquired pesticides from ChemService (West Chester, PA, USA). Nominal concentrations of pesticides (μg/L) were: 64 chlorpyrifos, 101 malathion, 171 terbufos, 91 aldicarb, 219 carbaryl, 209 carbofuran, 123 acetochlor, 127 alachlor, 105 metolachlor, 102 atrazine, 202 simazine, and 106 propazine. One hour after pesticides were applied, composite water samples were collected from mesocosms with the same pesticide treatment. These samples were shipped on ice to Mississippi State Chemical Laboratory to verify nominal concentrations. Measured concentrations of pesticides (μg/L) were: 60 chlorpyrifos, 105 malathion, 174 terbufos, 84 aldicarb, 203 carbaryl, 227 carbofuran, 139 acetochlor, 113 alachlor, 114 metolachlor, 117 atrazine, 180 simazine, and 129 propazine. The experimental design also included water and solvent (0.0001% acetone) controls (Fig. 1A). The experiment ran for four weeks, from June to July.

**Figure 1.**
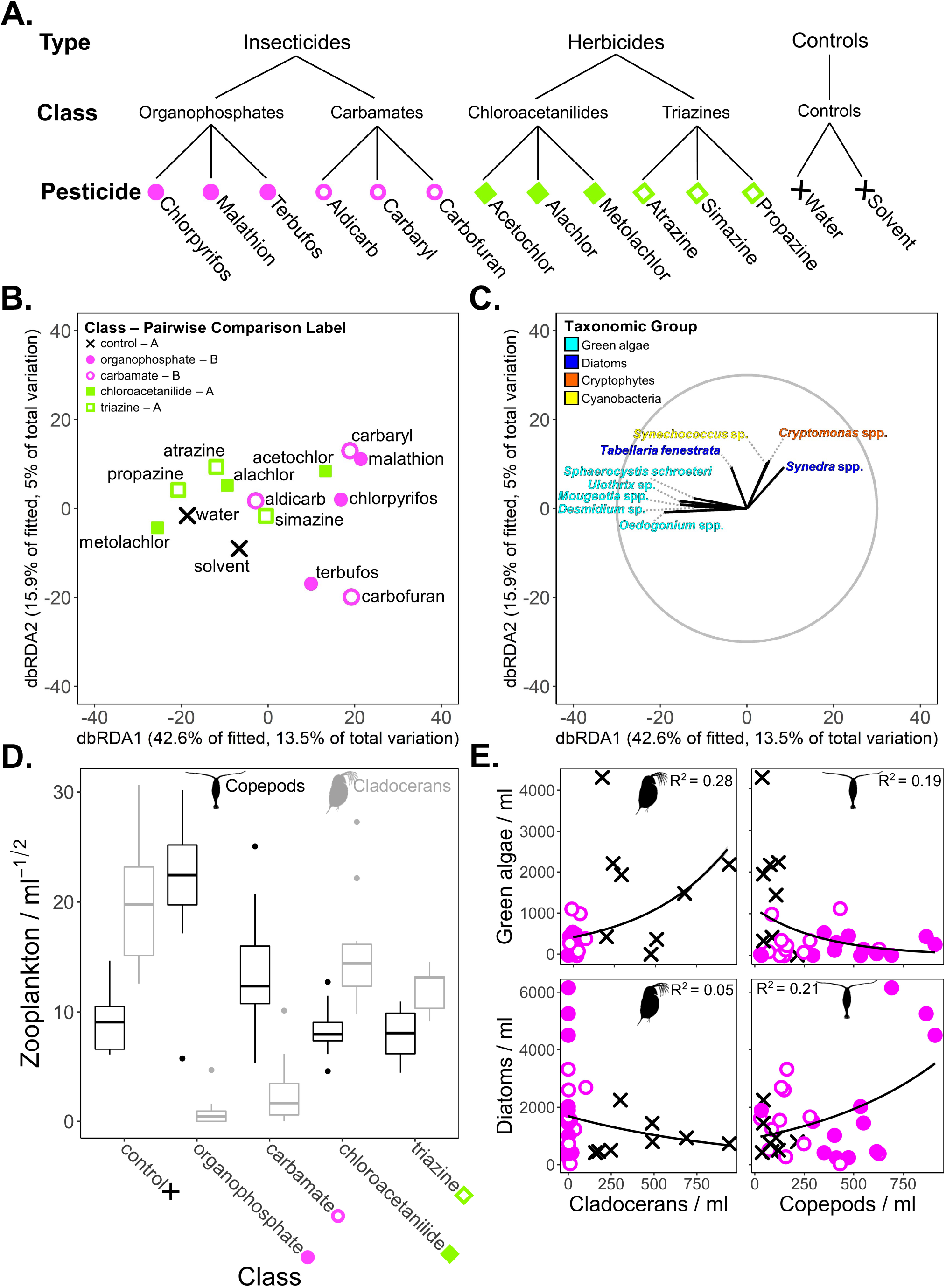
**A)** Experimental design highlighting the nested structure of pesticide treatments. Each treatment was replicated four times with mesocosm as the replicate. **B)** Distance-based redundancy analysis (dbRDA) plot of the phytoplankton community, showing differences among treatments by pesticide type. Points are the centroids of the 12 treatments. Treatments sharing the same letter are not different from each other in pairwise comparisons. **C)** Vector overlay of phytoplankton species or genera colored by broad taxonomic group, showing that insecticides were associated with a decrease in green algae and modest increase in diatoms. The gray circle corresponds to vector lengths that would have a correlation coefficient of one with a given axis. **D)** Cladoceran and copepod zooplankton densities in response to pesticide classes and the controls. Exposure to insecticides lead to copepods becoming more abundant compared to cladocerans. In contrast, herbicides reduced total zooplankton, but the relative amounts of cladocerans and copepods remained similar to the controls. **E)** Associations between densities of cladocerans or copepods and green algae or diatoms in insecticide and control treatments. Density of green algae was positively associated cladocerans and negatively associated with copepods, while diatoms were negatively associated with cladocerans and positively associated with copepods. In the plot of green algae and cladocerans, points have been jittered by 30 units in width and 50 units in height for ease of visualization. *R*^2^ values are McFadden pseudo-*R*^2^ values.

### Measurement of Experimental Responses

To characterize the algal community, we collected phytoplankton from the entire water column by inserting an upright PVC pipe, measuring 10 cm in diameter and 60 cm in height in the center of each mesocosm. A 1-liter subsample of the volume of water collected with the pipe was preserved with in a 1% solution of Lugol’s. Phytoplankton samples were collected in week four of the experiment. All samples were analyzed using the Utermöhl technique at 200-400 magnification (Lund *et al.* 1958). At least 400 natural units (colonies, filaments, and unicells) were enumerated using standard keys with taxonomy updated as necessary (Prescott 1962; Wehr & Kociolek 2015; Guiry & Guiry 2020). Densities per ml were calculated for a total of 74 genera and species of phytoplankton.

To characterize the zooplankton community, we collected zooplankton from the entire water column with a PVC pipe in the same manner as the phytoplankton samples; then, we capped the bottom and poured the water through 20 μm Nitex mesh. Two zooplankton samples were collected from each mesocosm, and these samples were combined and preserved in 70% ethanol. We counted and identified zooplankton to genera in 5 mL subsamples for each mesocosm with a zooplankton counting wheel (Wildlife Supply Company, Yulee, FL, USA) and a dissecting microscope. Zooplankton sampling occurred in week two of the experiment.

To measure total abundance of phytoplankton, we took 10 mL water samples, filtered phytoplankton onto glass fiber filters (under low vacuum pressure, <10 psi; Whatman EPM 2000, 0.3 μm, 47 mm), and measured chlorophyll-*a* concentrations of each sample. We used an organic extraction procedure with a 50:50 mixture of 90% acetone to DMSO and measured chlorophyll-*a* concentrations using a standard fluorometric technique (Carrick *et al.* 1993). Chlorophyll-*a* was measured from water samples taken in week two of the experiment.

To measure ecosystem respiration, we measured dissolved oxygen (DO) at dusk and dawn on subsequent days using hand-held meters (YSI, Yellow Springs, OH, USA) during weeks two and four of the experiment. Respiration was calculated as DO at dusk minus DO at dawn from the following day. Three previous manuscripts, which use the same design as the current manuscript, also describe this experimental design and methods in detail (Rohr *et al.* 2008; Rumschlag *et al.* 2019, 2020).

### Statistical Analyses

To evaluate the consistency of the effects of pesticides within type, class, and individual pesticide, we conducted a permutational multivariate analysis of variance (PERMANOVA, Table 1). This statistical model allowed us to attribute the variation explained in each pesticide level of organization (type, class, and individual pesticide), while accounting for the nested structure of our experimental design (Fig. 1A). The predictors were the following random categorical terms: type (insecticide, herbicide), class (carbamate, organophosphate, chloroacetanilide, triazine) nested with type, and pesticide (12 total) nested within class within type. No controls were included because they were not hierarchically nested (Fig. 1A). The multivariate response was a Bray Curtis similarity matrix based on a community matrix of square-root transformed densities of phytoplankton (abundance per ml) identified to genus or species at the end of the experiment. Across all treatments, 74 genera or species of phytoplankton were identified. The nested PERMANOVA used 9999 permutations and residuals under a reduced model. In addition, we evaluated pairwise differences between controls, carbamates, organophosphates, chloroacetanilides, and triazines with a multivariate comparison test using PERMANOVA (Fig. 1B). In this statistical model, 9999 unrestricted permutations of raw data were used. In both the nested and pairwise PERMANOVAs, Type III partial sums of squares were evaluated.

**Table 1.** Results of PERMANOVA models evaluating the effects of pesticides on the densities of algal communities. The multivariate response in the first model includes 74 species and genera. The multivariate response in the second model includes five broad taxonomic groups, including diatoms, green algae, cryptophytes, cyanobacteria, and euglenoids. *P* values were generated by Monte Carlo sampling, and those less than 0.05 are bolded. Variation explained is the proportion of the estimated component of variation for a given predictor relative to the model’s total variation.

| Endpoints and Source of Variation | *df* | Pseudo *F* | *P* | Variation Explained |
| --- | --- | --- | --- | --- |
| Algal community by species and genera |  |  |  |  |
| Type | 1 | 5.08 | **0.001** | 0.238 |
| Class(Type) | 2 | 0.68 | 0.845 | 0.000 |
| Pesticide(Class(Type)) | 8 | 1.17 | 0.175 | 0.134 |
| Residual | 34 |  |  | 0.628 |
| Algal community by taxonomic group |  |  |  |  |
| Type | 1 | 18.20 | **0.009** | 0.218 |
| Class(Type) | 2 | 0.12 | 0.971 | 0.000 |
| Pesticide(Class(Type)) | 8 | 1.77 | 0.052 | 0.242 |
| Residual | 34 |  |  | 0.539 |

To gauge how consistent the responses of phytoplankton community members are within broad taxonomic groups, we completed a second nested PERMANOVA on the algal community simplified to broad taxonomic groups (green algae, diatoms, cryptophytes, cyanobacteria, and euglenoids). (Table 1). This second nested PERMANOVA followed the same methods as the first, except that the multivariate response was based on densities of green algae, diatoms, cryptophytes, cyanobacteria, and euglenoids. This community matrix was generated by summing abundances of phytoplankton species and genera within their respective taxonomic groups. To determine how consistent the responses of phytoplankton were to pesticide exposures, we compared the relative amount of variation explained by the predictors and the residual variation in the two PERMANOVA models: the genera/species-level model and the model including the five taxonomic groups (Table 1).

To visualize 1) the consistency of the pesticide effects within type, class, and individual pesticide, 2) the consistency of responses of phytoplankton within broad taxonomic groups, and 3) how pesticides altered community composition of phytoplankton, we used a distance-based redundancy analysis (dbRDA, Fig. 1B, C), which is an ordination technique that constrains the community response matrix by environmental variables, in this case the pesticide treatments. The dbRDA was based on Bray-Curtis similarities of the square root transformed densities of the more than 70 genera or species of phytoplankton. The underlying categorical predictors were organophosphate, carbamate, chloroacetanilide, triazine, and control. In the dbRDA plot, we show the centroid values for the 14 experimental treatments. Both PERMANOVA models and the dbRDA were first executed using PERMANOVA+ for PRIMER version 7 (PRIMER-E Ltd, Plymouth, UK). Then, for ease of visualization of the dbRDA point and vector plots, data from PERMANOVA+ for PRIMER were exported, and plots were made using ‘*ggplol2*’ package in R.

Changes in zooplankton communities were evaluated using box-and-whisker plots (Fig. 1D). Namely, we examined the changes in the relative abundance of two major taxonomic groups of zooplankton, cladocerans and copepods, in response to classes of pesticides and the controls. To evaluate the associations between the densities of zooplankton and phytoplankton groups, we conducted four simple generalized linear regressions with Poisson distributions using the variation produced in the controls and the insecticide treatments (Fig. 1E). In these models, the response was either density of green algae or diatoms and the predictors were either cladocerans or copepods. For each model, we evaluated Type II sums of squares, a Wald χ^2^ test statistic, and a McFadden’s pseudo-R^2^.

To explore how exposure to pesticides altered abundance of broad taxonomic groups of phytoplankton, we examined box-and-whisker plots of densities of broad taxonomic groups of phytoplankton in response to classes of pesticides and the controls (Fig. 2A). In addition, we examined box-and-whisker plots of chlorophyll-*a*, a metric of total phytoplankton abundance (Fig. 2B). Since chlorophyll-*a* was measured soon after pesticide applications, we argue that chlorophyll-*a* measurements better reflect the immediate impacts of treatments on total abundance of phytoplankton compared to total phytoplankton density calculated from physical counts, which were measured at the end of the experiment.

**Figure 2.**
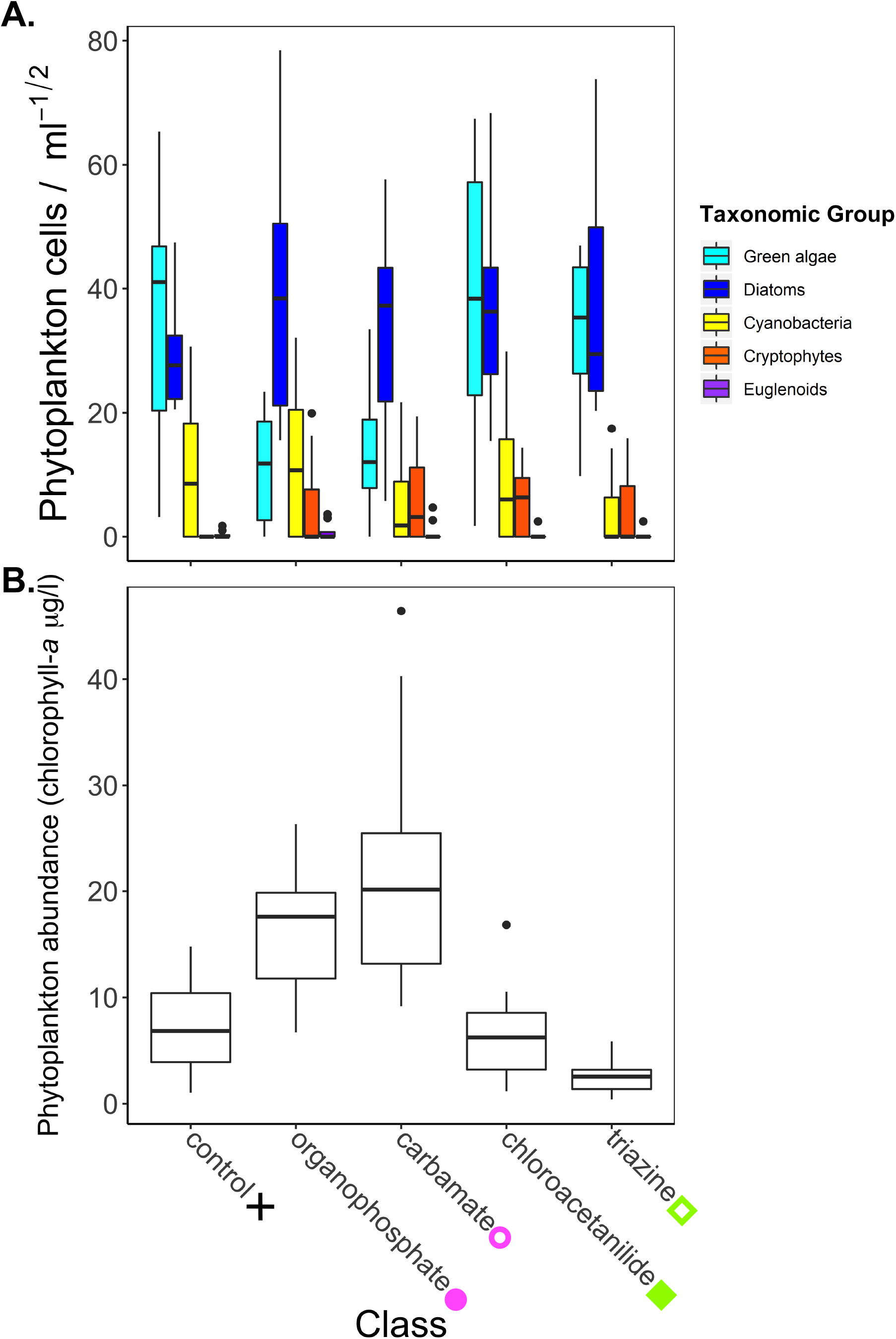
**A)** Densities of broad taxonomic groups of phytoplankton in response to pesticide classes and controls at the end of the experiment. Exposure to insecticides resulted in a decrease in green algae and an increase in the mean abundance of diatoms relative to controls. **B)** Total abundance of phytoplankton, as reflected by chlorophyll-*a* measurements, in response to pesticide classes and controls midway through the experiment. Insecticides were associated with increases in phytoplankton, while triazine herbicides were associated with decreases in phytoplankton relative to the control.

To examine how respiration of aquatic systems is altered by pesticides over time, we compared the effects of pesticides by class to controls on respiration at two- and four-weeks using log response ratios (Fig. 3A). For each time point, log response ratios of ecosystem respiration for individual mesocosms exposed to pesticides were calculated relative to the average of the control treatments (solvent and water controls) within a given spatial block. These calculations allow us to evaluate the magnitude of the difference in respiration of pesticide-treated mesocosms relative to the controls. We dropped observations in which DO at dawn was equal to DO at dusk, which generated an undefined value for the log response ratio (n = 2).

**Figure 3.**
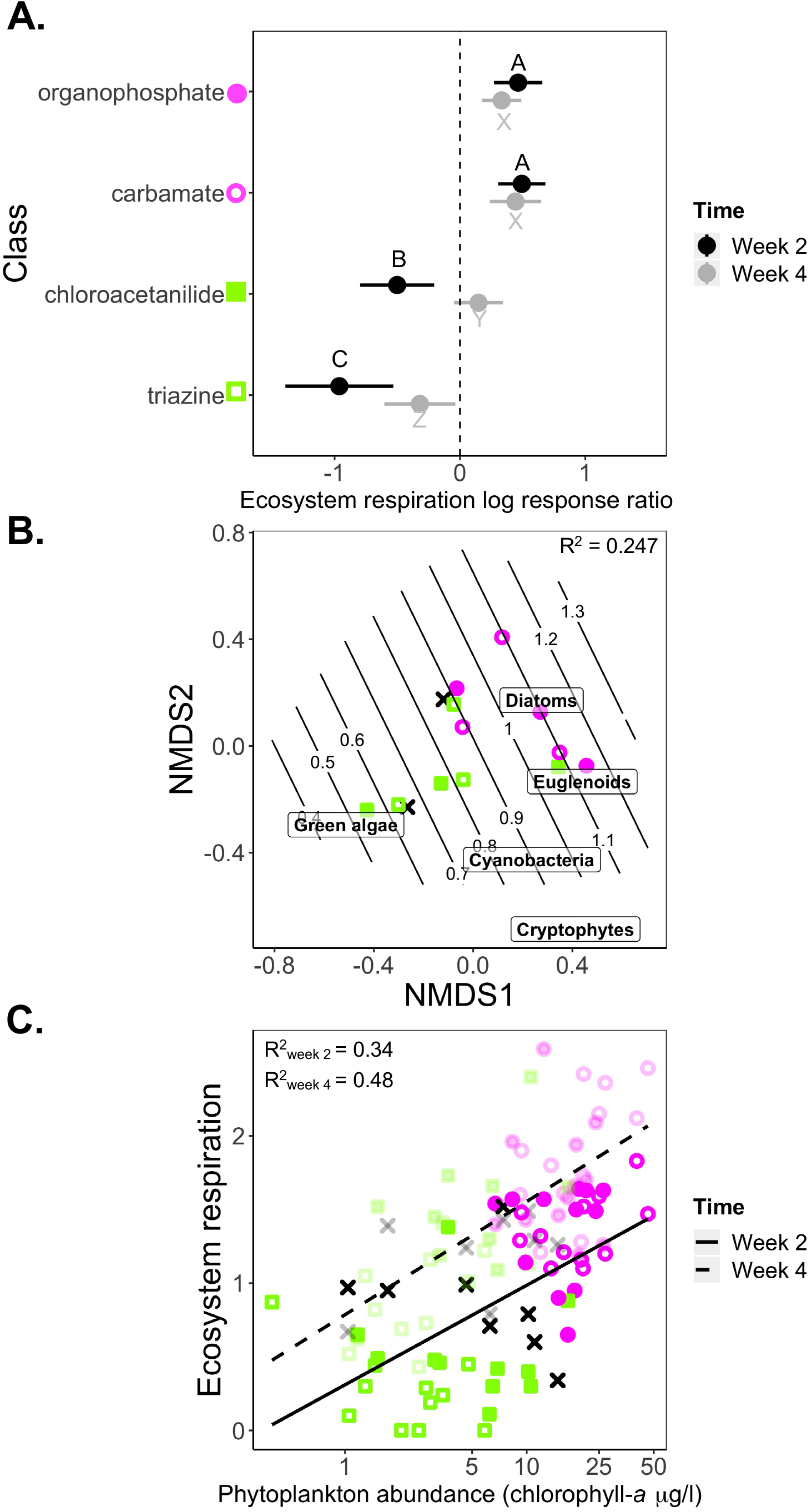
**A)** Log response ratios and 95% confidence intervals showing the effect of pesticides by class on ecosystem respiration at week two and four relative to the controls. Pairwise comparisons were conducted within time points. Classes sharing the same letter are not different from each other. Insecticides resulted in an increase in ecosystem respiration that persisted through week four. In contrast, herbicides were initially associated with a decrease in ecosystem respiration that did not persist to week four. **B)** NMDS of the phytoplankton community by broad taxonomic group (k = 3, stress = 0.10) and contour overlap of ecosystem respiration from week two. This plot shows that the increase in ecosystem respiration in insecticide treatments was associated with more diatoms and euglenoids and less green algae, relative to all other treatments. Herbicide treatments grouped with controls, suggesting herbicide-induced changes in algal community composition are not linked to ecosystem respiration. **C)** Regression plot showing a positive association between the abundance of phytoplankton, as measured by chlorophyll-*α*, and ecosystem respiration at weeks two and four suggesting that changes in the abundance of phytoplankton, as the result of herbicide and insecticide exposures, influenced ecosystem respiration. Associations at week two are shown with the solid line and opaque points. Associations at week four are shown with the dotted line and translucent points.

Next, to test how the community composition of phytoplankton was related to respiration measured at two weeks, we used non-metric multidimensional scaling (NMDS) ordination of broad taxonomic groups of phytoplankton densities based on Bray-Curtis similarities (‘metaMDS’ function, vegan package, Fig. 3B). Following NMDS ordination, we fit respiration from week two, using generalized additive models with the ‘ordisurf function (vegan package), which is a test of how well the ordination of the phytoplankton community predicts respiration (Fig. 3B). Finally, we assessed how the abundance of phytoplankton, as measured by chlorophyll-*a*, was related to respiration at two and four weeks using simple linear regressions (Fig. 3C). For the two models, the response was respiration measured at either two or four weeks, and the predictor was log-transformed chlorophyll-*a*. We evaluated Type II sums of squares. All analyses and plots, with the exception of the PERMANOVA models, were completed in R version 3.6.1. Preliminary analyses revealed no significant influence of the spatial block, so the spatial block was not included in the analyses.

## Results

### Pesticide types have consistent effects on broad taxonomic groups of phytoplankton

To evaluate the consistency of the effects of pesticides, we conducted a PERMANOVA that allowed us to attribute the variation explained in the phytoplankton community by pesticide type, class, and individual pesticide. For phytoplankton communities composed of more than 70 species and genera, pesticide type explained 24% of the variation in these communities (Table 1), which shows that the effects of pesticides on phytoplankton communities are generalizable to pesticide type.

To gauge the consistency in the responses of phytoplankton community members within broad taxonomic groups, we compared the genera/species-level PERMANOVA model to a model with the phytoplankton community simplified to broad taxonomic groups. Similar to the genera/species-level model, pesticide type explained 22% of the variation in the taxonomic groups (Table 1). Further, the residual variation in the model including broad taxonomic groups of phytoplankton was less than the genera/species-level model (54% versus 63% of residual variation, respectively, Table 1). Together, these results suggest that pesticides generally have similar effects on community members from the same broad taxonomic groups.

### Insecticides generate top-down effects on algal community composition and abundance which is associated with persistent increases in ecosystem respiration

We visualized the consistency in the effects of pesticides by type and the consistency in the responses of phytoplankton by broad taxonomic groups using dbRDA (Fig. 1B, C). The dbRDA showed that insecticide exposures significantly decreased green algae and modestly increased diatoms (Fig. 1B, C). These changes in community composition were likely the result of top-down effects of the zooplankton community. Insecticides had direct toxic effects on cladocerans, which lead to a competitive release of copepods (Fig. 1D). Across insecticides and controls, density of green algae was positively associated with cladocerans (*p* <.001, Wald χ^2^= 8495, pseudo-R^2^= 0.28, Fig. 1E top-left) and negatively associated with copepods (*p* <.001, Wald χ^2^= 4378, pseudo-R^2^= 0.19, Fig. 1E, top-right). At the same time, diatoms were negatively associated with cladocerans (*p* <.001, Wald χ^2^= 1455, pseudo-R^2^= 0.05, Fig 1E, bottom-left) and positively associated with copepods (*p* <.001, Wald χ^2^= 8329, pseudo-R^2^= 0.21, Fig.1E, bottom-right). In addition to changes in community composition, total abundance of phytoplankton, as indicated by chlorophyll-*a*, increased with insecticide exposure (Fig. 2B).

Comparisons of the effects of pesticides by class to controls using log-response ratios demonstrated that respiration of the entire aquatic community increased significantly in response to insecticide exposure, and this effect persisted throughout the experiment (Fig. 3A). The increase in ecosystem respiration was associated with both changes in community composition and total abundance of phytoplankton. For instance, the NMDS ordination of phytoplankton community composition and the vector overlay of ecosystem respiration at week two showed a significant association (*p* <.001, F_9,44_ = 1.93, R^2^= 0.25, Fig. 3B). The increase in ecosystem respiration in insecticide treatments was associated with more diatoms and euglenoids and less green algae relative to all other treatments (Fig. 3B). In addition, the greater total abundance of phytoplankton, represented by chlorophyll-*a*, in insecticide treatments was positively associated with ecosystem respiration in weeks two and four (week 2: *p* <.001, F_1,52_ = 26.88, R^2^= 0.34; week 4: *p* <.001, F_9,44_ = 48.34, R^2^= 0.48, Fig. 3C).

### Herbicides decrease phytoplankton abundance which is associated with short-term decreases in ecosystem respiration

In contrast to the effect of insecticides, the dbRDA and pairwise comparisons demonstrated no difference in community composition of phytoplankton between the herbicides and the controls (Fig. 1B, 2A). Instead, relative to controls, herbicide exposure resulted in a significant decrease in total abundance of phytoplankton, as indicated by chlorophyll-*a* (Fig. 2B). Log-response ratios showed that herbicides reduced ecosystem respiration, but these changes did not persist for the length of the experiment (Fig. 3A). In addition, the effects of herbicides on ecosystem respiration varied by class, with triazines having a greater magnitude of effect compared to chloroacetanilides (Fig. 3A). The NMDS ordination of the phytoplankton community composition and the vector overlay of ecosystem respiration at week two showed herbicides grouping with controls, suggesting no herbicide-induced changes in phytoplankton community composition could be linked to ecosystem respiration (Fig. 3B). Instead, the lower total abundance of phytoplankton (represented by chlorophyll-*a*) in herbicide treatments was associated with lower ecosystem respiration in weeks two and four (Fig. 3C).

## Discussion

Synthetic chemicals represent a globally widespread disturbance that threatens freshwater ecosystems. Yet, what remains largely unknown is how consistent the responses of a diversity of community members are to an array of contaminants and how these changes in communities correspond to alterations in the retention and release of carbon in ponds. We evaluated the effects of 12 pesticides, nested in four pesticide classes and two pesticide types on pond ecosystems, and included diverse phytoplankton and zooplankton communities. Our results demonstrate that: 1) the effects of pesticides were consistent within herbicide and insecticide types and within broad taxonomic groups of phytoplankton, 2) herbicides decreased total phytoplankton abundance but had no effect on phytoplankton composition, 3) by shifting dominant zooplankton from cladocerans to towards copepods, insecticides indirectly increased phytoplankton abundance and altered phytoplankton community composition via a top-down effect, and 4) herbicides lead to short-term decreases in ecosystem respiration that varied by herbicide classes, whereas insecticides lead to persistent increases in ecosystem respiration.

First, our results suggest that prediction of the staggering number of possible direct and indirect effects associated with freshwater communities being exposed to thousands of synthetic chemicals could be simplified to groups of chemicals with similar environmental targets and to broad groups of taxonomically related organisms. Given that only 0.36% of the more than 100 million unique chemicals currently in existence have gone through regulation by a federal agency (Gessner & Tlili 2016) and that freshwater systems are home to more than 125,000 described species (Dudgeon *et al.* 2006; Balian *et al.* 2008), society requires a risk assessment approach that can efficiently screen a vast number of chemicals against whole communities and ecosystems to accurately predict environmental safety; risk assessment using traditional approaches of testing a single compound against a single model organism could be extended to include tests of whole communities and ecosystems (Rohr *et al.* 2006, 2016; Clements & Rohr 2009). Simplifying prediction to groups of chemicals with similar environmental targets and responses of related organisms would improve efficiency for federal regulating bodies around the world and allow for more resources to be devoted to looking for exceptions to general patterns (Rohr *et al.* 2006, 2016; Clements & Rohr 2009).

The generalizable effects of insecticides and herbicides on algal and zooplankton communities that we found were consistent with previous studies. In the literature, insecticide exposure regularly reduce cladoceran zooplankton, which leads to a competitive release of copepods (Bridges & Boone 2003; Boone *et al.* 2005; Relyea & Diecks 2008; Relyea 2009; Hua & Relyea 2014). Additionally, these changes in the zooplankton community regularly have topdown effects on phytoplankton, consistent with the observed increase in total abundance of phytoplankton and a shift in the phytoplankton community observed in our study (Bridges & Boone 2003; Boone *et al.* 2005; Relyea & Diecks 2008; Relyea 2009; Hua & Relyea 2014). More specifically, we found increased density of copepods was negatively associated with the density of green algae and positively associated with diatom density. Total phytoplankton abundance might have increased because, compared to cladocerans, copepods are less efficient phytoplankton feeders and have broader diets encompassing non-algal food sources (Sommer & Sommer 2006). Changes in the phytoplankton community could have been the results of differences in feeding preferences of copepods versus cladocerans. Alternatively, insecticides may have indirectly altered the availability of nutrients for phytoplankton. Insecticides could have been directly toxic to insect predators, which in turn could have altered herbivory of snail and tadpoles and therefore, nutrient availability (Peacor & Werner 2000). Herbicides did not change phytoplankton composition but decreased total phytoplankton abundance; reduction in phytoplankton have been shown in other studies (Rohr & Crumrine 2005; Halstead *et al.* 2014, 2018). Similar to other studies, the total abundance of zooplankton in herbicide-treated ponds likely indirectly decreased because of the decrease in phytoplankton, a common food source for all zooplankton (Noack *et al.* 2003; Relyea 2009).

Few studies have attempted to evaluate how synthetic chemical disturbances in communities are associated with changes in ecosystem functions (Bernhardt *et al.* 2010; Rosi-Marshall & Royer 2012; but see McMahon *et al.* 2012; Rosi-Marshall *et al.* 2013; Halstead *et al.* 2014; Rumschlag *et al.* 2020). Our study found that insecticides led to persistent, and non-trivial, increases in ecosystem respiration, while herbicides lead to short-term decreases in ecosystem respiration that varied with pesticide classes. Alterations in the abundance and composition of phytoplankton caused by exposure to insecticides and herbicides could have been linked to changes in ecosystem in three ways, either individually or in combination. First, phytoplankton may have contributed directly to ecosystem respiration. So, decreased phytoplankton abundance in herbicide treatments might have resulted in less total respiration, while increased phytoplankton abundance and altered composition in insecticide treatments might have resulted in more total respiration. Second, the abundance of phytoplankton may have driven the abundance of microbial decomposers, which in turn could have contributed to ecosystem respiration. Finally, the abundance of phytoplankton may have led to more and/or larger secondary and/or tertiary consumers, which contributed to ecosystem respiration.

The variation in the duration and magnitude of changes in ecosystem respiration by pesticide type and class could be explained by the environmental persistence of the pesticides and the generation time of the organism to which the pesticide is directly toxic. For instance, ecosystem respiration in ponds exposed to herbicides might have recovered more quickly because herbicides were directly toxic to phytoplankton communities, whose generation times are upwards of a day (Laws 2013), and likely were able to rebound from the direct toxicity of herbicides. In contrast, ecosystem respiration in ponds exposed to insecticides might have been slower because direct toxicity affected zooplankton that have generation times of several weeks (Kalff 2002). In addition to ecosystem respiration varying by pesticide type, ecosystem respiration also varied by herbicide class. Triazine herbicides had a greater and more persistent negative effect on ecosystem respiration than chloroacetanilide herbicides, likely because triazines persist in the environment longer than chloroacetanilides (soil half-lives of 110-146 days versus 14-26 days respectively [Pesticide Acton Network Pesticide Database]).

Pesticide-induced alterations to respiration found in our study suggest that chemical contamination has the potential to alter the ecosystem metabolism in freshwater systems. While the influence of nutrient subsidies (nitrogen, phosphorus, and carbon) on metabolism in lakes and streams is well studied (Woodward *et al.* 2012; Stanley *et al.* 2016; Williamson *et al.* 2020), few studies have examined the influence of chemical contaminants, such as pesticides, pharmaceuticals (but see Rosi-Marshall *et al.* 2013; Robson *et al.* 2020), or heavy metals (but see Carlisle & Clements 2005), on components of the carbon cycle. In our study, while insecticides led to persistent increases in ecosystem respiration and a bloom in phytoplankton, insecticides could have led to carbon storage as biomass, release of atmospheric carbon, or no change in net primary productivity compared to controls. The outcome of the effect of insecticides on these patterns would depend on the size of the contribution of phytoplankton blooms to gross primary productivity. For instance, if gross primary productivity was greater or less than respiration, then carbon would have been stored as biomass or released, respectively. In contrast, we posit that herbicides could have led to a short-term release of atmospheric carbon because herbicides were associated with short-term decreases in respiration and phytoplankton abundance, which likely translated to reduced gross primary productivity. Future studies should investigate the effects of pesticides on the carbon cycle more holistically by directly evaluating changes to gross and net primary productivity and rates of carbon storage. These additional pieces of the carbon cycle would allow for the evaluation of how synthetic chemicals alter net fluxes and pools of carbon in aquatic systems. Runoff from spring application of pre-emergent herbicides on agricultural fields could possibly lead to short-term releases of atmospheric carbon in adjacent water bodies. Given the enormous amount of herbicides released in the environment annually (1.3 billion kg worldwide in 2012; Atwood & Paisley-Jones 2017) and that inland freshwater systems account for 0.6 billion tons of carbon storage (more than all carbon buried in oceanic sediments, Battin *et al.* 2009; Aufdenkampe *et al.* 2011), we postulate that herbicide use could have profound impacts on the global carbon cycle.

Given that the production and application of synthetic chemicals has been increasing exponentially for decades (Gessner & Tlili 2016), understanding the generalizable mechanisms by which synthetic chemicals can alter aquatic ecosystems is critical if our goal is efficiency in environmental risk assessment (Rohr & Crumrine 2005; Rohr *et al.* 2006, 2016). Our results support the hypothesis that predicting the effects of synthetic chemicals on complex, diverse freshwater systems can be simplified by generalizing patterns to groups of chemicals that share similar targets and to groups of organisms that are taxonomically related. To better understand, regulate, and protect human and ecological health, policy makers should look to generalizable patterns so that efficiency in risk assessment can become a priority, which will free resources to look for exceptions to general patterns.

## Acknowledgements

### Funding

Funding was provided by the National Institutes of Health (R01GM109499, R01TW010286-01), the National Science Foundation (EEID-1518681, EF-1241889, IOS-1754868, IOS-1651888), and the Department of Agriculture (NRI 2009-35102-0543).

### Competing interests

The authors declare no competing interests.

### Data and materials availability

All data are available at Figshare under the **accession number** XXXXXXX.

## Notes

### Competing Interest Statement

The authors have declared no competing interest.

